# *Nb*RD21 protease controls receptor kinase homeostasis in *Nicotiana benthamiana*

**DOI:** 10.64898/2026.04.22.720176

**Authors:** Alice Godson, Lauren Eddie, Mariana Schuster, Kaijie Zheng, Reka Toth, Yuge Li, Tianrun Li, Jie Huang, Farnusch Kaschani, Philippe V. Jutras, Jiorgos Kourelis, Markus Kaiser, Tolga Bozkurt, Renier A. L. van der Hoorn

## Abstract

RD21-like proteases are papain-like cysteine proteases with a C-terminal granulin domain that are abundant and ubiquitous in angiosperms and have often been implicated in immunity. We previously found that the activity of RD21 in *Nicotiana benthamiana* (*Nb*RD21) is suppressed during infection with *Pseudomonas syringae*. Here, we studied the role of *Nb*RD21 in immunity and proteome processing. *NbRD21* was disrupted by genome editing and *rd21* mutants were subjected to disease assays and shot-gun proteomics. Dipeptide substrate zLR-AMC was used in protease assays and agroinfiltration was used to transiently express *Nb*RD21 and candidate substrates. Genome edited lines lacking *Nb*RD21 develop normally but have drastically reduced zLRase activity and are significantly more susceptible to *P. syringae*. Shot gun proteomics revealed an increased accumulation of ∼20 diverse receptor-like kinases (RLKs) in untreated *rd21* knockout lines, but their transcript levels are unaltered when compared to wild-type plants. 35S-driven GFP-tagged RLKs accumulate more upon transient expression in *rd21* plants than in wild- type plants. These data indicate that *Nb*RD21 post-translationally controls RLK homeostasis, either by directly degrading RLKs, or indirectly by regulating endocytic RLK recycling.

## INTRODUCTION

Responsive-to-Desiccation-21 (RD21) -like proteases are ubiquitous in angiosperm plants. These papain-like cysteine proteases (PLCPs) are typified by having a C-terminal granulin domain and are amongst the most active and abundant PLCPs in vegetative tissues. RD21-like proteases accumulate in the vacuole, lytic vesicles, ER-bodies, and in the apoplast (reviewed by Huang & Van der Hoorn, 2025). Orthologs of Arabidopsis RD21 (At1g47128) are C14 in tomato, CP1A in maize, oryzain in rice and triticain in wheat. We previously also identified the granulin-containing *Nb*C14 in *N. benthamiana* (Kaschani et al., 2010; Bozkurt et al., 2011) but this protease belongs to PLCP subfamily-4 instead of subfamily-1 (Richau et al., 2012) and is therefore not an RD21 ortholog. *Nb*C14 has therefore been appropriately renamed into *Nb*CP14 (Paireder et al., 2016), since it is orthologous to *Nt*CP14, which is involved in developmental cell death during embryo development (Zhao et al., 2013).

Subfamily-1 RD21-like proteases are frequently associated with abiotic and biotic stresses. The *RD21* gene is abundantly transcribed in non-stressed plants, and is further induced upon drought and osmotic stress, hence the name responsive-to-dessiccation (Koizumi et al., 1993). RD21 proteases are also implicated in immunity and are often targeted by pathogen-secreted effectors that inhibit RD21 or trigger its degradation or mislocalisation (reviewed in Huang & Van der Hoorn, 2025). Tomato C14, for instance, is inhibited by cystatin-like EpiC inhibitors secreted by *Phytophthora infestans* (Kaschani et al., 2010) and by chagasin-like CIP1 produced by *Pseudomonas syringae* (Shindo et al., 2016), whilst the RD21 ortholog in maize (CP1) is inhibited by Pit2 (Misas-Villamil et al., 2019) and rice *Os*RD21 is inhibited by MoErs1 (Liu et al., 2024). Consistent with immune proteases being suppressed by pathogens, *rd21* mutant Arabidopsis lines are more susceptible to *Botrytis cinerea* (Shindo et al., 2012) and mutant wheat lines are more susceptible to wheat yellow mosaic virus (WYMV, Liu et al., 2023). However, how RD21-like proteases act in immunity has not yet been resolved.

We recently studied the suppression of apoplastic hydrolases of *Nicotiana benthamiana* (*Nb*) during infection by *Pseudomonas syringae* (*Ps*) pathovar *tomato* DC3000 (*Pto*DC3000) lacking Type-III effector HopQ1-1, which is pathogenic on *N. benthamiana* (Wei et al. 2007). Activity-based proteomics on apoplastic fluids revealed 60 apoplastic hydrolases that were suppressed upon infection (Sueldo et al., 2024). *Nb*RD21 (NbL08g19980) was one of the strongly suppressed hydrolases during infection. Here, we studied if *Nb*RD21 acts in immunity by taking advantage of the versatility of the *Nb/Ps* pathosystem.

## RESULTS

### Transient *Nb*RD21 overexpression increases protease activity and immunity

To investigate if *Nb*RD21 contributes to plant immunity by overcoming its suppression, we first transiently overexpressed *Nb*RD21 and its catalytic mutant (*Nb*RD21^C161A^) by agroinfiltration. Western blot analysis of leaves transiently overexpressing wild-type *Nb*RD21 with anti-C14 antibody (Kaschani et al., 2010) revealed a 36 kDa signal that represents intermediate *Nb*RD21, consisting of the catalytic domain and the granulin domain, and a 26 kDa mature *Nb*RD21, consisting of only the catalytic domain (**Figure 1A**). Transient expression of the *Nb*RD21^C161A^ mutant caused a high accumulation of the intermediate isoform and a low accumulation of the mature isoform (**Figure 1A**). These data are consistent with the catalytic mutant of Arabidopsis *At*RD21 expressed by agroinfiltration (Gu et al., 2012) and demonstrate that *Nb*RD21 also removes the granulin domain autocatalytically. The prodomain of *Nb*RD21^C161A^, however, is still removed, consistent with the removal of the prodomain of *At*RD21 by other proteases (Yamada et al., 2001).

**Figure 1.**
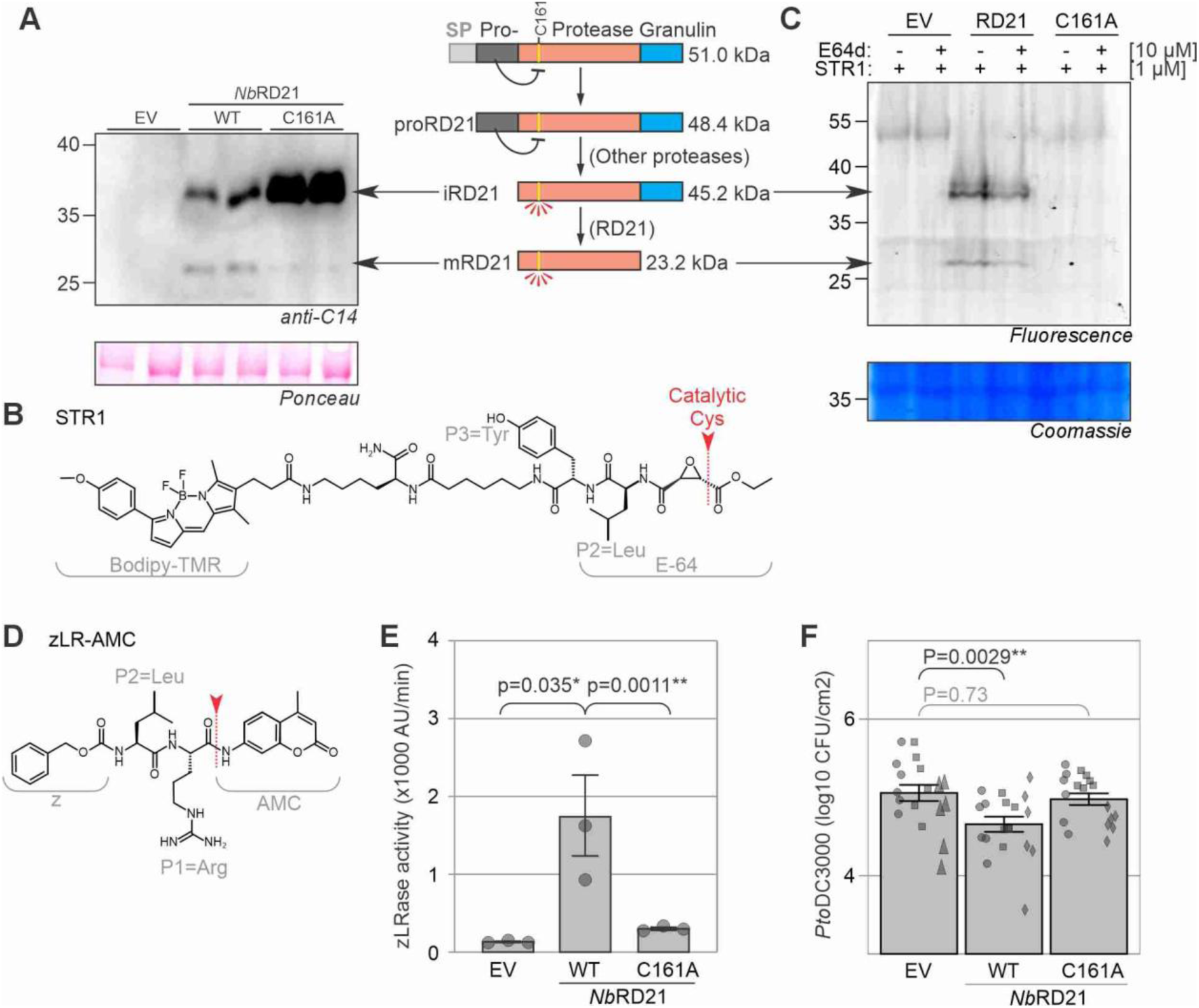
Activity-dependent immunity upon *Nb*RD21 overexpression **(A)** The *Nb*RD21 catalytic mutant accumulates in agroinfiltrated *N. benthamiana* leaves. *Nb*RD21 and its catalytic mutant (C161A) and the empty-vector (EV) control were co-expressed with silencing inhibitor P19 by agroinfiltration and total extracts were isolated at 5dpi, separated by SDS-PAGE and analysed by western blotting with α-C14 antibody and Ponceau staining. **(B)** Molecular structure of activity-based probe STR1 probe for PLCPs. STR1 consists of the E-64 warhead linked to a Bodipy-TMR fluorophore. **(C)** Both intermediate and mature *Nb*RD21 are labelled with STR1. *Nb*RD21 and its catalytic mutant (C161A) and the empty-vector (EV) control were co-expressed with silencing inhibitor P19 by agroinfiltration and total extracts were isolated at 5dpi, and preincubated with 10 μM E-64d for 1 hr and then labeled with 1 μM STR1 for 3 hrs. Samples were separated on protein gel and detected by in-gel fluorescence scanning with Cy2 settings at PMT700. **(D)** Molecular structure of the PLCP substrate zLR-AMC. **(E)** *Nb*RD21 has zLRase activity. Total extracts from plants transiently expressing wildtype or the C161A mutant of *Nb*RD21 or an EV control was isolated at 5 dpi and incubated with 40 μM zLR-AMC in the presence of 2.5 mM DTT and sodium acetate pH 5. Fluorescence was measured at 380ex/460em at 15 minutes at room temperature in a plate reader. Error bars represent mean ± SE of n=3 replicates. Statistical significance was calculated with the Student’s *t*-test; *, P<0.05. **(F)** *Nb*RD21 requires catalytic activity for its immune function. *Nb*RD21, its catalytic site C161A mutant, and the EV control were transiently co-expressed by agroinfiltration in 4-week-old *N. benthamiana* with silencing inhibitor P19. Two days later, the same leaves were infiltrated with 10^6^ CFU/mL *Pto*DC3000. Bacterial growth in colony-forming units (CFU) was measured at 3 days after PtoDC3000 inoculation. Bars represent SE of n=6 replicates of three experiments. P-values were calculated by two-way ANOVA followed by post-hoc comparison using the Dunnett test. **, P<0.01.

To demonstrate which isoform is active, we custom-synthesized a fluorescent activity-based probe STR1, consisting of the covalent PLCP inhibitor E-64 coupled to a Bodipy fluorophore (**Figure 1B**, Huang et al., 2026). This probe is identical to the previously used MV201 (Richau et al., 2012), except that STR1 lacks the azide minitag on the Bodipy fluorophore. Labeling of total extracts of leaves transiently overexpressing *Nb*RD21 with 2 µM STR1, followed by separation on SDS-PAGE and detection by fluorescence scanning revealed that both the intermediate and mature isoforms of *Nb*RD21 are fluorescently labeled (**Figure 1C**), indicating that they are both active proteases. Preincubation with 10 µM E-64 suppresses labeling. The intermediate and mature isoforms of *Nb*RD21^C161A^, however, are not labeled, consistent with being inactive and lacking the catalytic cysteine that would be labeled by STR1 (**Figure 1C**).

To further demonstrate that the *Nb*RD21 is active and *Nb*RD21^C161A^ not, we used commercially available zLR-AMC, a Leu-Arg dipeptide linked to the 7-amino-4-methyl coumarin (AMC) fluorophore (**Figure 1D**). Incubation of total leaf extracts with zLR-AMC revealed that *Nb*RD21 overexpression strongly increased zLRase activity (**Figure 1E**). By contrast, extracts containing *Nb*RD21^C161A^ had zLRase activity similar to the empty vector (EV) control (**Figure 1E**).

Having established that *Nb*RD21 overexpression increases STR1 labelling and zLRase activity, and *Nb*RD21^C161A^ not, we next monitored growth of wild-type (WT) *Pto*DC3000 in these agroinfiltrated tissues through the ‘agromonas’ assay (Buscaill et al., 2021). At two days after agroinfiltration, leaves were infiltrated with wild-type *Pto*DC3000 and the colony forming units (CFU) were determined three days later. *Nb*RD21 overexpression caused a significant reduction in bacterial growth, in contrast to *Nb*RD21^C161A^ which displayed similar susceptibility as the EV control (**Figure 1F**). Thus, *Nb*RD21 overexpression increases immunity to WT *Pto*DC3000 in an activity-dependent manner, consistent with being an immune protease.

We next tested bacterial growth of the Δ*hopQ1-1* mutant of *Pto*DC3000, which is pathogenic on *N. benthamiana* because it lacks HopQ1-1, the Type-III effector that is recognized in *N. benthamiana* by Roq1 (Wei et al., 2007). Remarkably, bacterial growth of *Pto*DC3000(*ΔhQ*) is indistinguishable between leaves overexpressing *Nb*RD21 and the EV control (Supplemental **Figure S1A**). We also tested bacterial growth of pv. *tabaci* 6605 (*Pta*6605) and pv. *syringae* B728a (*Psy*B728a) and found that bacterial growth was indistinguishable between leaves overexpressing *Nb*RD21 and the EV control (Supplemental **Figures S1B and S1C**). Thus, *Nb*RD21 suppresses growth of WT *Pto*DC3000, which is recognized by Roq1, but not of *P. syringae* strains that cause disease.

### Gene edited *rd21* mutants have reduced protease activity and increased susceptibility

To investigate if *Nb*RD21 is also required for immunity, we disrupted the open reading frame of *Nb*RD21 (NbL08g19980) in *N. benthamiana* with CRISPR/Cas9 genome editing. We selected two independent homozygous lines with truncated gene products: the *rd21-1* line has a 4 base pair deletion, and the *rd21-2* line has a 1 base pair insertion (**Figure 2A**). We also sequenced two potential off-site targets of the used sgRNAs but found that two RD21-like genes (NbL04g16800 and NbL17g14770) were unaltered in the *rd21* mutants (Supplemental **Figure S2**).

**Figure 2.**
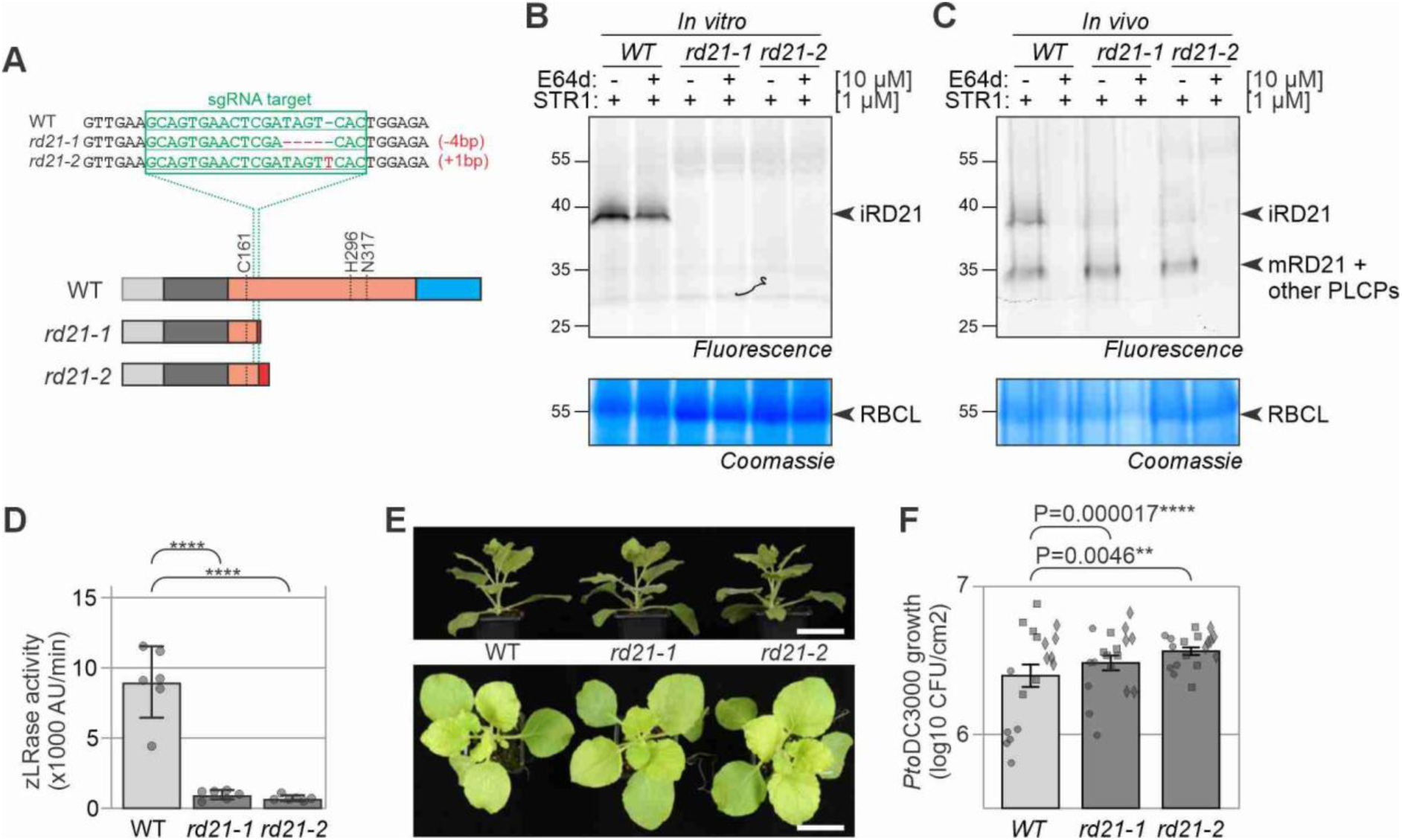
*rd21* mutant plants have reduced zLRase activity and increased susceptibility. **(A)** Two *rd21* mutants contain frameshift-inducing indels in the sgRNA targeted region. Mutants *rd21-1* and *rd21-2* carry a 4 bp deletion and a 1 bp insertion, respectively. **(B**) Reduced *in vitro* PLCP labeling in both *rd21* mutants. Leaves from 4wk old plants were injected with 10 μM E-64d or Mock and incubated in the dark 1 hr and then extracts were made and labeled with 1 μM STR1 for 2 hrs. Proteins were separated by SDS-PAGE and scanned for fluorescence or stained with Coomassie. **(C)** Reduced *in vivo* PLCP labeling in both *rd21* mutants. Leaves of 4 wk old plants were injected with 10 μM E-64d or Mock and incubated in the dark for 1 hr and then injected with 1 μM STR1 and incubated for 2 hrs. Leaf extract were generated, separated by SDS-PAGE and scanned for fluorescence and stained with Coomassie. **(D)** Both *rd21* knockouts have significantly reduced zLRase activity. Protease activity was measured in total leaf extracts using 40 µM zLR-AMC. Error bars represent SE of n=6 replicates. P-values were determined using one-way ANOVA and Dunnett’s multiple comparisons test. **(E)** Both *rd21* mutants have no growth or developmental phenotype. Shown are representative photos of four-week-old plants. Scale bar, 7 cm. **(F)** *Pto*DC3000 growth is increased in *rd21* knockout lines. Leaves of wildtype (WT) and *rd21* knockout plants were infiltrated with wildtype *Pto*DC3000 at OD_600_=0.0001. Bacterial growth in colony-forming units (CFU) was measured at 3 dpi. Bars represent SE of n=6 replicates of three experiments. P-values were calculated by two-way ANOVA followed by post-hoc comparison using the Dunnett test.

Activity-based labeling of total extracts with STR1 demonstrated that the fluorescent 36 kDa signal is absent in the mutants (**Figure 2B**), indicating that this signal is caused by intermediate *Nb*RD21*. In vivo* labeling by infiltrating STR1 into leaves also revealed that the 36 kDa signal is absent in both *rd21* mutants (**Figure 2C**). The zLRase activity is also significantly reduced in total leaf extracts of *rd21* mutants when compared to the WT control (**Figure 2D**), demonstrating that *Nb*RD21 caused the major zLRase activity in total extracts. Despite lacking *Nb*RD21, the *rd21* mutant plants have no macroscopic developmental or growth phenotype (**Figure 2E**). Infection assays revealed that *rd21* plants are slightly but significantly more susceptible to wild-type *Pto*DC3000 (**Figure 2F**), consistent with reduced growth of *Pto*DC3000 upon *Nb*RD21 overexpression (**Figure 1F**).

### Proteomic analysis of *rd21* mutants

To identify changes in the proteome of *rd21* mutants, we performed shotgun proteomics on total leaf extracts of both *rd21* lines and the WT control and annotated the spectra to the LAB360 proteome (Ranawaka et al., 2023). A total of 2,779 proteins of *N. benthamiana* were detected and PCA analysis revealed that the *rd21* mutants are distinct from WT samples (Supplemental **Figure S3**). We therefore combined these *rd21* samples for a statistical analysis to identify differentially accumulating proteins. We identified 15 proteins that accumulate significantly less in the *rd21* mutants, including *Nb*RD21 itself (**Figures 3A and 3B**, Supplemental **Table S1**). Interestingly, four Rab GTPases also accumulate significantly less in the *rd21* mutants: Rab1 (NbL10g13430), Rab2 (NbL19g08250), Rab11 (NbL14g13650) and Rab8a (NbL08g02890) (**Figure 3C**).

**Figure 3.**
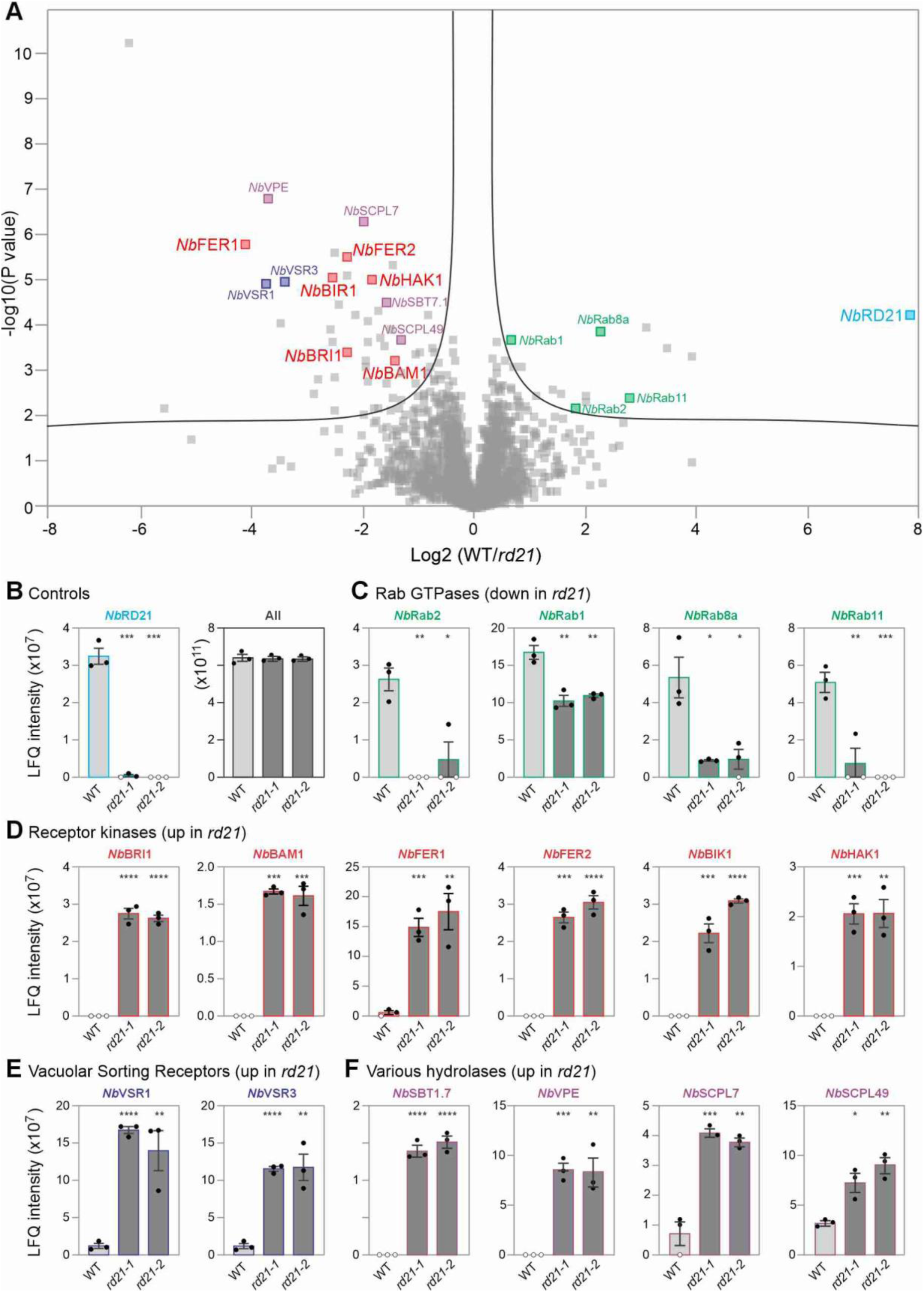
Differential protein accumulation in *rd21* knockout plants. **(A)** Volcano plot of MS data showing log_2_ of the fold-change in LFQ-intensity between wildtype and both *rd21* knockout lines against the significance (-log_10_(P-value)) of this difference. All data points above the black line have significant P values (t-test, FDR=0.05, S0=0.1). **(B-F)** Bar charts showing raw LFQ intensities in wildtype (WT) and both knockout (*rd21-1* and *rd21-2*) plants. Open dots indicate the absence of a detectable MS signal. All in (B) is the sum of the intensities of all detected peptides of *Nicotiana benthamiana* proteins. Error bars represent SE of n=3 samples. Student’s *t*-test compared to WT values: *, p<0.05; **, p<0.01; ***, p<0.001, ****, p<0.0001.

Amongst the 49 proteins that accumulate significantly more in both *rd21* mutants (Supplemental **Table S1**) are six receptor-like kinases (RLKs) of which four having extracellular leucine-rich repeats (LRRs): BAM1 (Barely-Any-Meristem-1, NbL19g02860), BRI1 (Brassinosteroid Insensitive-1, NbL15g00030), BIR1 (BAK1-Interacting-RLK-1, NbL09g13380), HAK1 (Herbivory-associated Kinase-1, NbL05g07580), and two having extracellular malectin-like domains: FER1 (Feronia-1, NbL05g07500) and FER2 (Feronia-2, NbL19g15210) (**Figure 3D**). Other proteins that accumulate significantly more in the *rd21* mutants are two vacuolar sorting receptors (VSR1 (NbL08g14990) and VSR3 (NbL17g22450), **Figure 3E**) and various hydrolases including a vacuolar processing enzyme (VPE, NbL08g18900), subtilisin SBT1.7 (NbL16g14350), and two serine carboxy peptidases-like proteins (SCPL49 (NbL05g22050) and SCPL7 (NbL18g14840), **Figure 3F** and Supplemental **Table S1**).

To test if increased RLK levels in *rd21* plants is associated with higher transcript levels of the corresponding RLK-encoding genes, we performed quantitative RT-PCR on these plants with primers specific for *FER1, BRI1, BAM1* and *HAK1*, using *eIF4A* as a control. This RT-qPCR experiment demonstrated that transcript levels of these four RLKs are not different in the *rd21* mutants (Supplemental **Figure S4**), indicating that the higher RLK accumulation is a post-translational process.

To investigate the higher accumulation of RLKs in *rd21* mutants further, we searched for all detected RLKs and evaluated their differential LFQ intensity. Besides the six RLKs discussed above, we identified 16 additional RLKs in the proteomics dataset. Three RLKs accumulated to similar LFQ intensities in *rd21* mutants when compared to WT plants (Supplemental **Figure S5**), but the remaining 13 RLKs seem to accumulate more in the *rd21* mutants, although for 10 RLKs this is not statistically significant because they were not detected in sufficient samples (Supplemental **Figure S6**). RLKs that accumulate more in *rd21* plants as well as the RLKs that have that tendency carry very different ectodomains and belong to different RLK subfamilies assigned in the metaRLK database (Zhang et al., 2026, Supplemental **Figure S7**). Likewise, the three RLKs that accumulate to similar levels are unrelated: two RLKs have different LRR ectodomains and the other RLK carries a lectin-G ectodomain (Supplemental **Figure S7**). In conclusion, although protein accumulation is unchanged for three RLKs, most RLKs accumulate higher in *rd21* plants, indicating that increased protein levels are common for diverse RLKs in *rd21* mutants.

### Transient expression confirms increased RLK accumulation in *rd21* mutants

To confirm the increased accumulation of RLKs in the *rd21* mutants, we transiently expressed *Nb*BAM1-GFP and *Nb*FER2-GFP driven by a constitutive 35S promoter in the *rd21* mutants and WT control plants and monitored their accumulation by measuring GFP fluorescence. GFP fluorescence was detected for *Nb*BAM1-GFP but not for *Nb*FER2-GFP (**Figure 4A**). *Nb*BAM1-GFP fluorescence was significantly increased in both *rd21* mutants (**Figures 4A and 4B**), consistent with the higher accumulation of *Nb*BAM1 in *rd21* plants in the proteomics experiment. Confocal imaging showed that GFP fluorescence of *Nb*BAM1-GFP accumulates in the plasma membrane and in endosome-like vesicles but this subcellular localization seems not different between WT and *rd21* mutants (**Figure 4C**).

**Figure 4.**
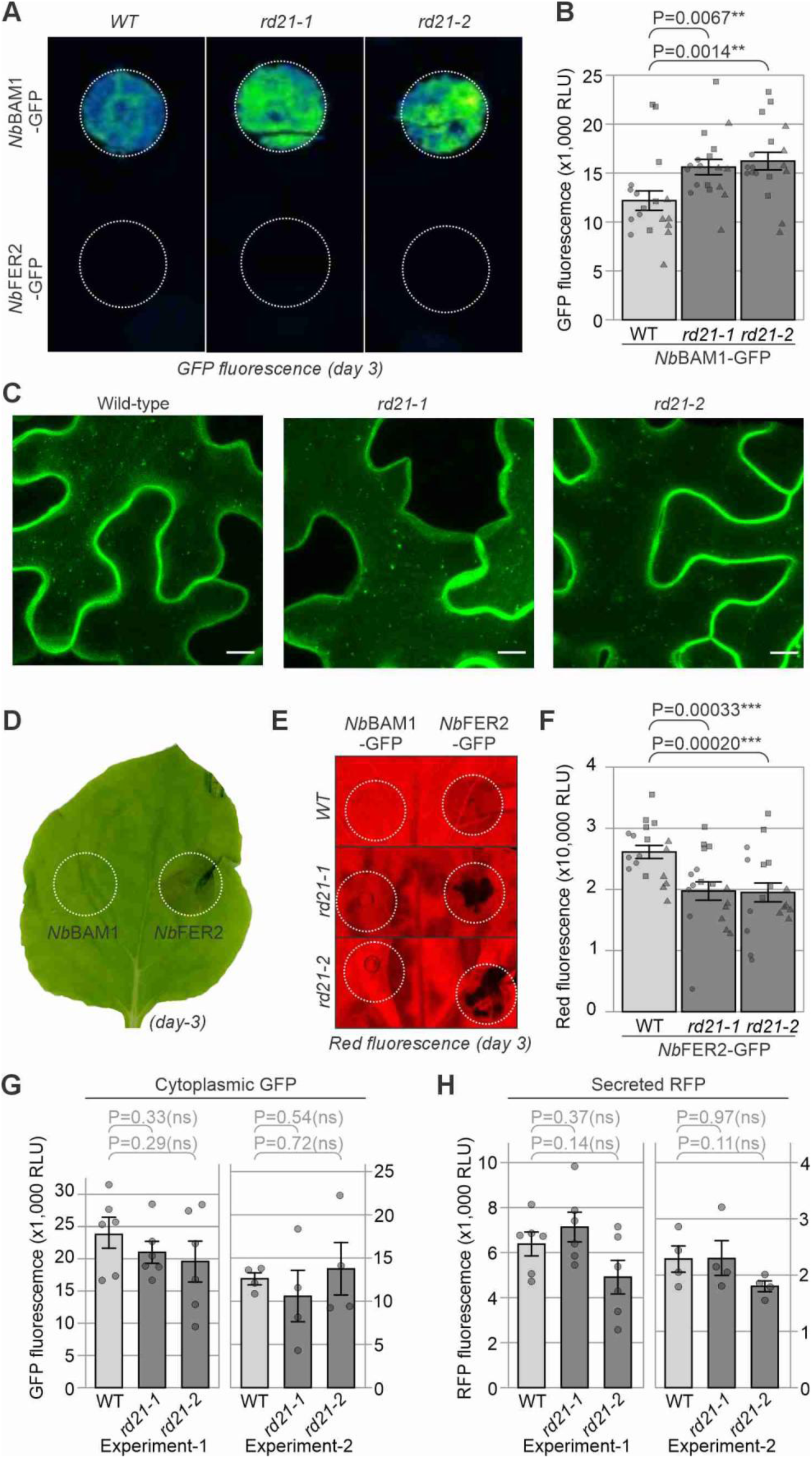
***Nb*BAM1 and *Nb*FER2 levels are increased in *rd21* plants. (A)** GFP fluorescence upon transient expression of *Nb*BAM1-GFP and *Nb*FER2-GFP. Shown are agroinfiltrated regions expressing *Nb*BAM1-GFP (top) and *Nb*FER2-GFP (bottom). Scale bar, 1cm. **(B)** Quantified GFP fluorescence upon transient expression of *Nb*BAM1-GFP. Bars represent SE of n=6 replicates of three experiments. P-values were calculated by two-way ANOVA followed by post-hoc comparison using the Dunnett test. **, P<0.01. **(C)** Z-projection of confocal images of transiently expressed *Nb*BAM1-GFP in various genotypes at day-4. Scale bar, 10 µm. **(D)** Cell death triggered by transient expression of *Nb*FER2-GFP at day 3. **(E)** Red autofluorescence upon transient expression of *Nb*BAM1-GFP and *Nb*FER2-GFP. **(F)** Quantified red autofluorescence upon transient expression of *Nb*FER2-GFP. (A-F) *Nb*BAM1-GFP and *Nb*FER2-GFP were transiently expressed in WT and *rd21* plants and images were taken 3 days later. Bars represent SE of n=6 replicates over three experiments. P-values were calculated by two-way ANOVA followed by post-hoc comparison using the Dunnett test. **(G)** Quantified GFP fluorescence upon transient expression of cytoplasmic GFP (cGFP) is unaltered in *rd21* plants. **(H)** Quantified RFP fluorescence upon transient expression of secreted RFP (sRFP) is unaltered in *rd21* plants. (G-H) cGFP and sRFP were transiently expressed in WT and *rd21* plants and images were taken three days later. Bars represent SE of n=6 and n=4 plants of two experiments, respectively. P-values were calculated by Student’s t-test.

Transient expression of *Nb*FER2-GFP, however, did not produce sufficient GFP fluorescence (**Figure 4A**), but triggered cell death instead (**Figure 4D**). We therefore measured the reduction of leaf autofluorescence in the red spectrum as a quantitative readout for cell death (Xi et al., 2021). These measurements demonstrated that *Nb*FER2-GFP significantly reduces red autofluorescence in both *rd21* mutants when compared to the WT control plants (**Figure 4E**). Since cell death is a functional readout of *Nb*FER2-GFP, these data are consistent with the increased accumulation of this receptor in *rd21* plants.

To test if higher transient expression in *rd21* mutants is not caused by increased Agrobacterium susceptibility, we expressed cytoplasmic GFP (cGFP) and secreted RFP (sRFP) (Dodds et al., 2025) in WT and *rd21* plants and monitored their expression by measuring fluorescence. This experiment revealed that transient GFP expression causes fluorescence that is not significantly different in the *rd21* mutant compared to WT plants (**Figure 4G**). Likewise, also transient expression of secreted RFP caused fluorescence that is similar between WT and *rd21* plants (**Figure 4H**). Thus, the increased levels of BAM1-GFP fluorescence and FER2-GFP-induced cell death cannot be caused by increased transient expression in *rd21* mutants but involves a post-translational process that affects specific proteins.

## DISCUSSION

Protease activity assays established that *Nb*RD21 is responsible for most zLRase activity in leaf extracts. *Nb*RD21 overexpression suppresses growth of *Pto*DC3000, and this requires *Nb*RD21 activity. However, *Nb*RD21 overexpression did not suppress bacterial growth of virulent *P. syringae* strains. Conversely, *rd21* plants are significantly more susceptible to *Pto*DC3000. Proteomics uncovered that *rd21* mutants accumulate more hydrolases, vacuolar sorting receptors and receptor kinases, whilst four Rab proteins accumulate less in *rd21* plants. Further proteome studies indicated that most but not all RLKs accumulate in *rd21* plants and RT-PCR experiments indicated that this is not due to higher transcript levels of RLK-encoding genes. Transient expression of two of these RLKs indicate that they indeed accumulate more in *rd21* mutants. This higher accumulation is not caused by general improved transient expression in *rd21* plants because similar fluorescence levels were produced upon transient expression of cytoplasmic GFP or secreted RFP.

The observation that *Nb*RD21 overexpression only caused phenotypes with avirulent *Pto*DC3000 and not with the virulent *Pto*DC3000(*ΔhQ*), *Pta*6605 or *Psy*B728a is intriguing. The dependence of the disease phenotype in *Pto*DC3000 on the Type-III effector HopQ1-1 might suggest that HopQ1-1 is a substrate of *Nb*RD21, but this is unlikely, as HopQ1-1 is a cytonuclear effector (Li et al., 2013), whereas RD21 is mostly in the endomembrane system (Lampl et al., 2013; Ondzighi et al., 2008). Rather, we believe that the phenotype can be explained by the fact that HopQ1-1 triggers the effector-triggered immune (ETI) response that includes the release of the vacuolar content into the apoplast upon the fusion of the vacuolar and plasma membranes (Hatsugai et al., 2009). During ETI, *Pto*DC3000 will be exposed to large quantities of *Nb*RD21 released from the vacuole and this may directly or indirectly reduce bacterial growth. The absence of disease phenotypes with virulent strains might be explained by the fact that *Nb*RD21 concentrations in the apoplast are lower than the vacuole, and these *Nb*RD21 levels might be sufficiently suppressed during infection by CIP1 and other PLCP inhibitors (Shindo et al., 2016; Sueldo et al., 2024).

The *rd21* mutants showed a drastically reduced zLRase activity in total extracts, indicating that *Nb*RD21 is responsible for this activity. This zLRase activity probably comes from *Nb*RD21 directly because zLR-AMC is a good substrate for purified *Nb*RD21 *in vitro* (Paireder et al., 2017; named *Nb*CysP6 there). Indeed, zLRase activity is also strongly increased upon overexpressing *Nb*RD21 but not *Nb*RD21^C161A^. The decreased RbcL degradation during activity-based labeling of leaf extracts is consistent with the fact that RD21 is a major protease activity in *N. benthamiana*. In fact, RbcL degradation in Arabidopsis leaf extracts was even stronger in the presence of SDS, which activates latent RD21 (Gu et al., 2012). The diversity of substrates is consistent with *Nb*RD21 being a general protease, rather than a specific processing enzyme. Indeed, purified *Nb*RD21 can completely degrade various IgG antibodies *in vitro* and Proteomics Identification of Cleavage Sites (PICS) revealed that *Nb*RD21 has a broad substrate range, only preferring hydrophobic residues at the P2 position (Paireder et al., 2017). Despite these strong biochemical phenotypes, *rd21* mutants grow indistinguishably from wild-type *N. benthamiana*, which is similar to the absence of growth phenotypes in *rd21* mutants of *Arabidopsis thaliana* (Shindo et al., 2012).

Our analysis of leaf proteomes, however, revealed several proteins accumulate differentially in *rd21* plants. Perseus analysis of label-free quantification (LFQ) values revealed 49 proteins accumulating more in the *rd21* mutants, which could be direct and indirect *Nb*RD21 substrates, or result from mechanisms attempting to compensate for *Nb*RD21 loss. However, none of the hydrolases accumulating in the *rd21* mutants are predicted to replace the broad protease specificity of *Nb*RD21. Rather, we detect the accumulation of specific processing enzymes such as vacuolar processing enzyme (VPE), which cleaves after Asn(N) and Asp(D) residues (Yamada et al., 2020), and putative phytaspase SBT1.7, which seems to cleave after Asp(D) residues (Buscaill et al., 2024). Interestingly, VPEs and SBT1.7 might accumulate in response to reduced *Nb*RD21 activity. Prodomain removal of RD21 proteases is catalyzed by other proteases that are present in leaf extracts (Yamada et al., 2001). Although VPEs were proposed to be responsible because of Asp/Asn residues at the prodomain junction (Yamada et al., 2001), quadruple *vpe* mutants showed unaltered RD21 activation (Gu et al., 2012), indicating that other proteases might act redundantly with VPEs in RD21 activation. Interestingly, SBT1.7 is a phytaspase that cleaves after Asp(D) residues and might act redundantly with VPE in *Nb*RD21 activation. Thus, both VPE and SBT1.7 might be upregulated in an attempt to activate *Nb*RD21 in the *rd21* mutant. This activation by Asp-specific proteases is similar to the activation of papain-like Rcr3 protease of tomato by subtilase P69B cleaving at Asp(D) residues after the prodomain (Paulus et al., 2020). Like Rcr3, *Nb*RD21 also carries an Asp(D) at this position.

Proteome analysis revealed an increased accumulation of six RLKs in *rd21* plants. Further analysis of the proteomics data indicated that another 13 RLKs accumulate higher in *rd21* plants, whilst three RLKs accumulate equally well in *rd21* plants. These RLKs carry various ectodomains and belong to different subfamilies. RT-PCR analysis showed that the increased accumulation of FER1, BRI1, BAM1 and HAK1 is not associated with increased expression of genes encoding these RLKs.

These data suggest that *Nb*RD21 is involved in RLK homeostasis, either by directly degrading these proteins, or by regulating their protein levels otherwise. Indeed, the higher accumulation of transiently expressed *Nb*BAM1-GFP confirms the increased RLK accumulation. The fact that *Nb*BAM1-GFP is expressed from a constitutive 35S promoter and still differentially accumulates in *rd21* mutants implies that increased *Nb*BAM1 accumulation is not caused by differential gene expression but by post-translational regulation. Likewise, although the *NbBAM1* gene carries an intron, the *35S:BAM1-GFP* construct does not carry this intron, excluding the possibility that differential accumulation is caused by differential splicing. The absence of detectable GFP fluorescence with transiently expressed *Nb*FER2-GFP is probably caused by the cell death that is triggered by *Nb*FER2 expression, which is associated with pleiotropic phenotypes described for Feronia mutants in Arabidopsis (Malivert & Hamant, 2023). Cell death might only be triggered at high FER2 levels reached upon agroinfiltration because *rd21* plants have no phenotypes despite increased FER2 levels. Although the cause of this cell death is yet unclear, this cell death response is a functional readout of *Nb*FER2 activity and the increased cell death in the *rd21* mutants therefore implies that also *Nb*FER2-GFP accumulates to higher levels in the *rd21* mutant.

*Nb*RD21 might decrease RLK receptor levels by degrading them directly, or indirectly, by regulating RLK homeostasis, either before delivery to the plasma membrane, or after. Receptors that are not folded in the ER are selected by ER quality control (ERQC) and degraded through ER-associated protein degradation (ERAD). It is possible that *Nb*RD21 facilitates the degradation of misfolded receptors. However, *Nb*RD21 can also act in receptor-mediated endocytosis (RME), where receptors are sorted at the plasma membrane (PM) into vesicles which travel through the endosomal system and are either recycled back to the PM, or taken to the vacuole for degradation (Claus et al., 2018). A role of *Nb*RD21 in RME is supported by the increased accumulation of vacuolar sorting receptors VSR1 and VSR3 in the *rd21* mutants, and the reduced levels of Rab proteins involved in vesicle fusion in the *rd21* mutants. Furthermore, *At*RD21 also accumulates in the trans-Golgi network (TGN) and in lytic vesicles (Ondzighi et al., 2008) and cysteine protease inhibitor E-64 caused the accumulation of endosomes in starving BY-2 tobacco cell cultures (Yamada et al., 2005), suggesting that a PLCP like RD21 might facilitate endosomal trafficking to the vacuole. We did indeed detect *Nb*BAM1-GFP in vesicles but its subcellular localization is not altered in the *rd21* mutants.

The absence of developmental phenotypes and disease phenotypes with virulent *P. syringae* strains is remarkable, given the increased accumulation of ∼20 diverse RLKs in noninfected *rd21* plants. There are several explanations for this. First, BIR1 is a negative regulator of RLKs (Liu et al., 2016, Liu et al., 2023) and its increased accumulation might cancel out increased RLK activities. Second and similarly, the effect of increased RLK activity may be cancelled out by the reduced immunity described in *rd21* mutants (reviewed in Huang & Van der Hoorn, 2025). Third, increased RLK activities might have been cancelled out by Type-III effectors of *P. syringae*, many of which effectively suppress RLK activities (Block & Alfano, 2011). But besides cancelling out increased RLK activity, it is also possible that the increased RLK protein levels represent a larger proportion of RLKs that are no longer functional and are destined for degradation, originating either from the ER or the PM. *Nb*RD21 might facilitate the removal of these nonfunctional RLKs, either directly, or indirectly.

Taken together, we have established important new tools in elucidating the functions of RD21-like proteases in plants and used these to discover their involvement in receptor homeostasis in plants. These *rd21* mutants and other materials and protocols will be instrumental for future research into receptor kinases and the roles of RD21 proteases.

## MATERIALS & METHODS

### Plants

*N. benthamiana* was grown in a growth chamber at room temperature with a 16-hour light cycle and 60% humidity and watered on alternating days.

### Molecular cloning

Used primers and plasmids are summarized in Supplemental **Table S2**. and Supplemental **Table S3**, respectively. *Nb*RD21^C161A^ was created by site-directed mutagenesis using pPB034 (Sueldo et al., 2024) as template. Overlapping primers incorporating the desired mutations were designed and the fragments amplified by PCR using Q5 Polymerase (New England Biolabs). DpnI was used to digest the methylated template. To generate binary constructs expressing C-terminally tagged RLKs, the Cf-4 sequence of pJK227 was replaced by the coding sequences of *Nb*FER2 and *Nb*BAM1 amplified from cDNA obtained from *N. benthamiana* leaf tissue via Gibson assembly and transformed into *Escherichia coli* DH10β. Sanger sequencing was used to verify the constructs before transformation into *Agrobacterium tumefaciens* GV3101 (pMP90) and selection for kanamycin and gentamicin resistance.

### Total extract (TE) isolation

Leaf discs (1 cm diameter) were punched from leaves and flash-frozen in liquid nitrogen. Leaf tissue was ground to a powder with a plastic pestle and resuspended in PBS (1:3 w:v). Samples were centrifuged at 14,000 x *g* for 10 minutes at 4^°^C and either used immediately or stored at -20^°^C.

### Western blot analysis

Proteins separated by SDS-PAGE were transferred to polyvinylidene difluoride (PVDF) membranes using the Trans-blot Turbo transfer system (Biorad). Membranes were blocked with 5% milk in PBS-T (PBS pH 6.1, 0.1% Tween-20). Membranes were rinsed in PBS-T before addition of primary antibody, and between addition of primary antibody (Kaschani et al., 2010) and secondary antibody. Membranes were finally washed in PBS. Antibodies were incubated with agitation for at least 1 hour at room temperature or overnight at 4°C. Chemiluminescence was measured using an ImageQuant LAS 4000.

### Genome editing

Cas9 vector pJK309, containing an *in planta* NPTII selection cassette, 2×35S-driven Cas9 module, and a multiple cloning site, was generated by combining pICSL4723, pICH47751 (Addgene plasmid #48002), pICH41766 (Addgene plasmid #48018) (Weber et al., 2011), pICSL11024 (Addgene plasmid #51144), pICH47742::2×35S-5’UTR-hCas9(STOP)-NOST (Belhaj et al., 2013; Addgene plasmid #49771) in a BsaI GoldenGate reaction. Two independent sgRNA sequences targeting NbL08g19980 (Supplemental **Table S2**) were cloned into the Cas9 vector pJK309 to produce pPJ64. Agrobacterium cultures were grown in LB media (kanamycin 50 μg/mL and gentamicin 10 μg/mL) overnight at 28°C. The next day, cultures were centrifuged at 4000 x *g* for 10 minutes at room temperature and resuspended in agroinfiltration buffer (10 mM MgCl_2_, 10 mM MES, 150 μM acetosyringone) to a final OD_600_ of 0.1. 4-week-old *N. benthamiana* plants were infiltrated and three days later, leaf discs were punched from infiltrated leaves, incubated in distilled water for one hour and soaked in 10% bleach for ten minutes. Leaf discs were incubated on shooting media without antibiotics for three days before transferring onto media containing kanamycin. Shoots were transferred to fresh shooting media every 1-2 weeks until transplantation into rooting media from week 6. Once roots were established, plantlets were potted and grown in the growth chamber. Seeds of heterozygous primary transformants were sown and genotyped.

### Genotyping

Genomic DNA was extracted from three plants of each genotype by crushing a leaf disc in extraction solution (1:10 GOWARDS,TEBY). Specific PCR products were generated using the primers described in Supplemental **Table S2**, designed to amplify the sgRNA-targeted regions. PCR products were separated on 1% agarose gel at 90V to check successful amplification. DNA was purified by ethanol precipitation, the DNA concentration measured using a nanodrop spectrophotometer, and samples sent for sequencing.

### zLRase assay

Cysteine protease activity was measured using the fluorogenic substrate Z-Leu-Arg-7-amino-4-methylcoumarin (zLR-AMC, R&D Systems). TE was mixed with 80 µM zLR-AMC in a black 96-well plate in the presence of 10 mM TCEP and 50 mM sodium acetate pH 5 at a final reaction volume of 100 μL. Fluorescence at 380 nm excitation / 460 nm emission was recorded in a plate reader (Tecan Infinite M200) at 20 second intervals for 10-60 minutes. Activity was calculated by change in fluorescence over time. P-values were determined by t-test or one-way ANOVA with Dunnett correction.

### Protease activity profiling

Activity-based labeling of PLCPs was performed as described previously (Richau et al., 2012), but now with STR1, a fluorescent E-64 synthesised by WuXi AppTec. TE samples were incubated with 1 μM STR1 in the presence of 50 mM sodium acetate pH 6.0 and 5 mM DTT in the dark for 3 hours at room temperature with agitation. Samples were precipitated with ice-cold acetone and resuspended in 2X SDS gel loading buffer (200 mM Tris-Cl pH 6.8, 400 mM DTT, 8% SDS, 0.4% bromophenol blue, 40% glycerol) before heating at 95°C for 5 mins. The proteins were separated on 12% acrylamide gels at 100V for 2.5 hours. The labelled proteins were detected by imaging the gel using the Typhoon scanner using Cy3 settings. Coomassie stain used to check equal protein loading.

### Disease assays

Bacterial growth assays were performed as described previously (Buscaill et al., 2021). *Pto*DC3000(ΔhQ) was grown in LB media at 28°C for 24 hours, centrifuged at 3,500 x *g* for 10 mins at room temperature and resuspended in water at OD_600_=0.0001. Six plants per genotype and three leaves per plant were syringe-infiltrated. Three days later, three 1 cm diameter leaf discs were taken per plant, surface sterilised with 15% hydrogen peroxide and washed in MilliQ water. Leaf discs were placed into Eppendorf tubes with 3 mm diameter metal beads and 1 mL MilliQ water and lysed in tissue lyser at 30 hertz per second for 5 mins. Leaf samples plated on LB agar plates containing cetrimide (10 μg/mL), fusidic acid (10 μg/mL) and cefaloridine (50 μg/mL) in a 10-fold serial dilution, and incubated at 28°C. Two days later, colonies were counted and converted to colony forming units (CFU) per cm^2^ of leaf surface. P-values were calculated using two-way ANOVA including an interaction factor and post-hoc Dunnett’s test to compare bacterial growth in each *rd21* mutant to wildtype plants, accounting for variation resulting from different replicate experiments.

### Proteome analysis

TE was isolated from wildtype and *rd21* mutant *N. benthamiana* plants in n=3 replicates. Protein concentration was measured using the DC assay (Bio-Rad). TE samples contained 200 µg protein in 100 µL. Halt Protease Inhibitor Cocktail (Thermo Fisher Scientific 78429) was added to samples at 1:100 to prevent proteolytic degradation. Samples were carbamidomethylated by reduction with 5 mM DTT for 30 minutes at 37°C followed by alkylation with 20 mM 2-chloroacetamide for 30 minutes at room temperature. Proteins were precipitated using methanol and chloroform (Wessel and Flügge, 1984), pellets were air-dried and proteins were eluted by sonicating in 6 M urea. Proteins were digested with trypsin/LysC at 1:40 enzyme:proteome for three hours at 37°C. Samples were then diluted in 50 mM Tris-HCl pH 8 (until <1 M urea) and incubated overnight at 37°C with shaking. The pH of the samples was adjusted to <3 with heptafluorobutyric acid (HFBA) and 100 µg protein was desalted on stage tips (Bond Elut OMIX C18, Agilent A57003100) first with 50% acetonitrile, then with 1% and 0.1% HFBA. The next day, peptides were eluted in 0.1% formic acid and 75% acetonitrile before being dried for 2 hours (SpeedVac).

### LC-MS/MS analysis

Each sample was analysed on an Orbitrap Fusion Lumos instrument (Thermo) that was coupled to an EASY-nLC 1200 liquid chromatography (LC) system (Thermo). The LC was operated in the one-column mode. The analytical column was a fused silica capillary (75 µm × 44 cm) with an integrated fritted emitter (15 µm; CoAnn Technologies) packed in-house with Reprosil-Pur 120 C18-AQ 1.9 µm resin (Dr. Maisch). The analytical column was encased by a column oven (Sonation) and attached to a nanospray flex ion source (Thermo). The column oven temperature was adjusted to 50 °C during data acquisition. The LC was equipped with two mobile phases: solvent A (0.1% formic acid, FA, in water) and solvent B (0.1% FA, 19.9% water and 80% acetonitrile, ACN). All solvents were of UPLC grade (Honeywell). Peptides were directly loaded onto the analytical column with a maximum flow rate that would not exceed the set pressure limit of 980 bar (usually around 0.6 – 1.0 µL/min). Peptides were subsequently separated on the analytical column by running a gradient of solvent A and solvent B.

### Peptide and Protein Identification using MaxQuant

RAW spectra were submitted to an Andromeda (Cox et al., 2011) search in MaxQuant (v2.5.2.0.) using the default settings (Cox & Mann 2008). Label-free quantification and match-between-runs was activated (Cox et al., 2014). The MS/MS spectra data were searched against the *N. benthamiana* (NbLab360.v103.gff3.CDS.fasta.AA.fasta, 45730 entries). All searches included a contaminants database search (as implemented in MaxQuant, 245 entries). The contaminants database contains known MS contaminants and was included to estimate the level of contamination. Andromeda searches allowed oxidation of methionine residues (16 Da) and acetylation of the protein N-terminus (42 Da) as dynamic modification. Carbamidomethylation on Cysteine (57 Da) was set as static modification. Enzyme specificity was set to “Trypsin/P specific”. The instrument type in Andromeda searches was set to Orbitrap and the precursor mass tolerance was set to ±20 ppm (first search) and ±4.5 ppm (main search). The MS/MS match tolerance was set to ±0.5 Da. The peptide spectrum match FDR and the protein FDR were set to 0.01 (based on target-decoy approach). Minimum peptide length was 8 amino acids and maximum amino acid length 25. For protein quantification unique and razor peptides were allowed. Modified peptides were allowed for quantification. The minimum score for modified peptides was 40. Label-free protein quantification was switched on, and unique and razor peptides were considered for quantification with a minimum ratio count of 2. Retention times were recalibrated based on the built-in nonlinear time-rescaling algorithm. MS/MS identifications were transferred between LC-MS/MS runs with the “match between runs” option in which the maximal match time window was set to 0.7 min and the alignment time window set to 20 min. The quantification is based on the “value at maximum” of the extracted ion current. At least two quantitation events were required for a quantifiable protein. Further analysis and filtering of the results was done in Perseus v1.6.10.0. (Tyanova et al., 2016). Comparison of protein group quantities (relative quantification) between different MS runs is based solely on the LFQ’s as calculated by MaxQuant, MaxLFQ algorithm (Cox et al., 2014).

### Proteomics analysis

MS data were searched against the LAB360 proteome database (Ranawaka et al., 2023) and a contaminants database (implemented in MaxQuant (version 2.0.2.0, Cox et al., 2011)). Data analysis, filtering and statistics of the MaxQuant output received from ACE was completed using the Perseus computer software (version 2.0.3.0, Tyanova et al., 2016). Missing values were imputed in order to calculate a volcano plot (wildtype – knockout). Categorical groups were compared by performing a Student’s t test (FDR=0.05; S0=0.1, 250 randomisations). The volcano plot was annotated in Perseus and edited in Inkscape. Significant hits were functionally annotated using information from the following webpages: BLAST, InterPro, Pfam, UniProt, and TAIR. Principal component analysis (PCA) was carried out using TBtools-II (Chen et al., 2023).

### Detection of RLK-GFP proteins

For quantifying GFP fluorescence, six plants per genotype were agroinfiltrated with *Nb*BAM1 and *Nb*FER2. Leaves were collected at 3 days-post infiltration and scanned for fluorescence with an Amersham Typhoon using filters Cy2 (GFP) and Cy5 (red fluorescence).

### RT-PCR

To measure the expression of PRRs in WT *N. benthamiana* and *rd21* mutants, similar age *N. benthamiana* leaves (∼100 mg) were harvested for Vazyme FastPure Universal Plant Total RNA Isolation Kit (#RC411-01) based RNA extraction, following manufacturer’s protocol. Next, cDNA synthesis was performed using SuperScript IV First Strand Synthesis System (#18091050) kit. Primers used in RT-qPCR are provided in Supplemental **Table S2**. NbeIF4A-14 (NbL17g05310.1) was used as internal reference gene. qPCR reactions were performed on StepOnePlus Real-Time PCR System (#15360337) using PowerUP SYBR Green Master Mix (#4368577) in a MicroAmp Fast Optical 96-Well Reaction Plate with Barcode (#4346906), following the manufacturer’s protocol. Relative gene expression was determined using the 2−ΔΔCt method.

### Transient cGFP/sRFP expression

To examine the expression of cGFP and sRFP in the WT and *rd21* mutants, at least four plants were agroinfiltrated. At three days-post infiltration, fluorescence signals were scanned using an Amersham Typhoon scanner with Cy2 (GFP) and Cy3 (RFP) filters.

### Confocal Microscopy

Transiently expressed *Nb*BAM1-GFP proteins were visualized in abaxial epidermal cells of wild-type and *rd21* mutant *N. benthamiana* leaves using a Zeiss LSM 880 confocal microscope with a C-Apochromat 40x/1.2 W Korr FCS M27 water-immersive objective 4 days post infiltration. GFP fluorescence was excited with 2% laser power at 488 nm and emission detected with a PMT detector at 493 to 598 nm. For z-stacks 16-bit images were captured at a slice interval of 1.31 μm and Z-projections made using Fiji image analysis software.

## Data availability

The mass spectrometry proteomics data for the on-bead digestions have been deposited to the ProteomeXchange Consortium via the PRIDE (Vizcaíno et al., 2016) partner repository (https://www.ebi.ac.uk/pride/archive/) with the dataset identifier PXD063276.

## ACKNOWLEDGEMENTS

We thank Ursula Pyzio for excellent plant care and Sarah Rodgers, Caroline O’Brien and Patricia Bowman for technical support.

## Author contributions

RH conceived the project; AG, LE, MS, and KZ performed most experiments with help of YL, TL and JH; PJ generated the *rd21* mutant, RT performed microscopy and TL performed RT-PCR and RLK classification; TB and JK provided materials and JH, FK and MK performed proteomic analysis; RH wrote the manuscript with input from all authors.

## SUPPLEMENTAL FIGURES and TABLES

**Figure S1.**
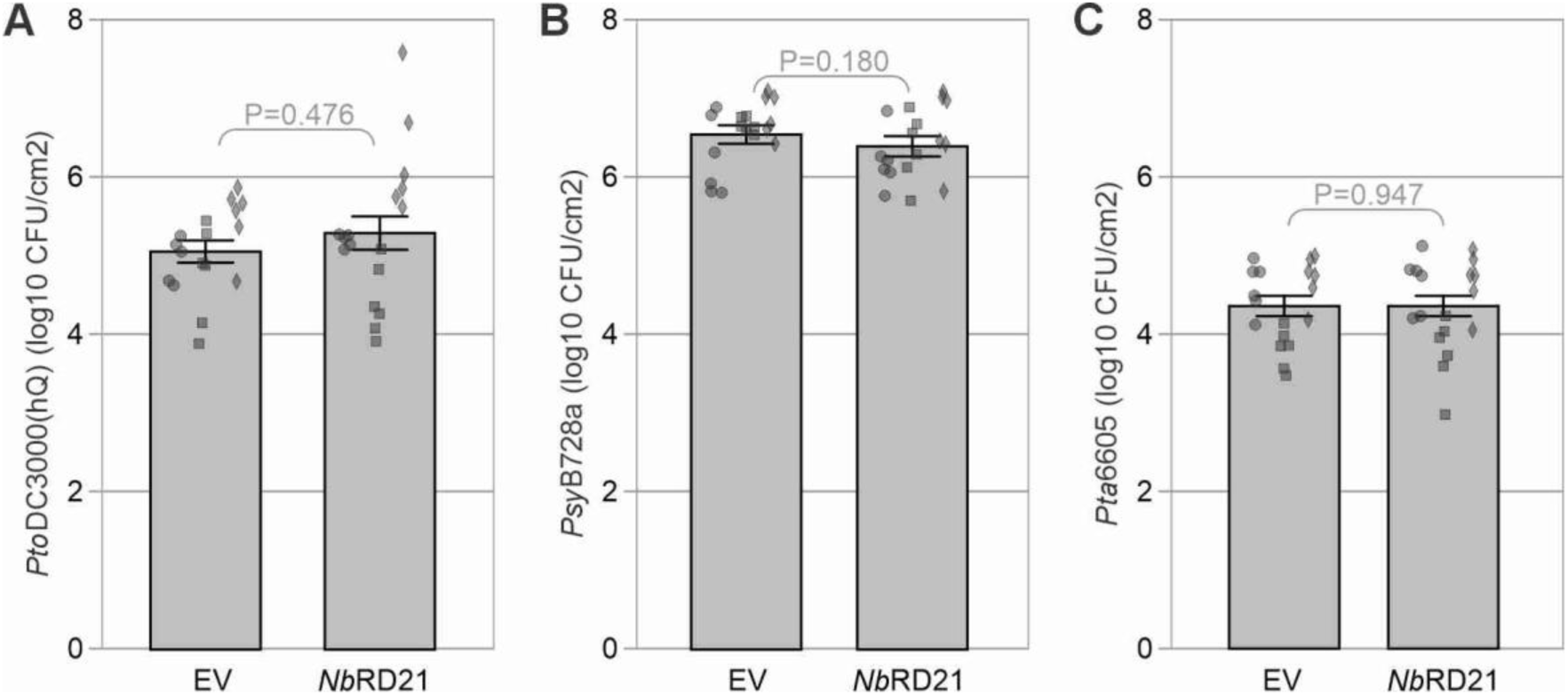
No differential bacterial growth in *rd21* mutants for virulent *P. syringae* strains. *Nb*RD21 and the EV control were transiently co-expressed by agroinfiltration in 4-week-old *N. benthamiana* with silencing inhibitor P19. Two days later, the same leaves were infiltrated with 10^6^ CFU/mL *P. syringae*. Bacterial growth in colony-forming units (CFU) was measured at 3 days after *Pto*DC3000 inoculation. Bars represent SE of n=6 replicates of three experiments. P-values were calculated by two-way ANOVA followed by post-hoc comparison using the Dunnett test. **, P<0.01.

**Figure S2.**
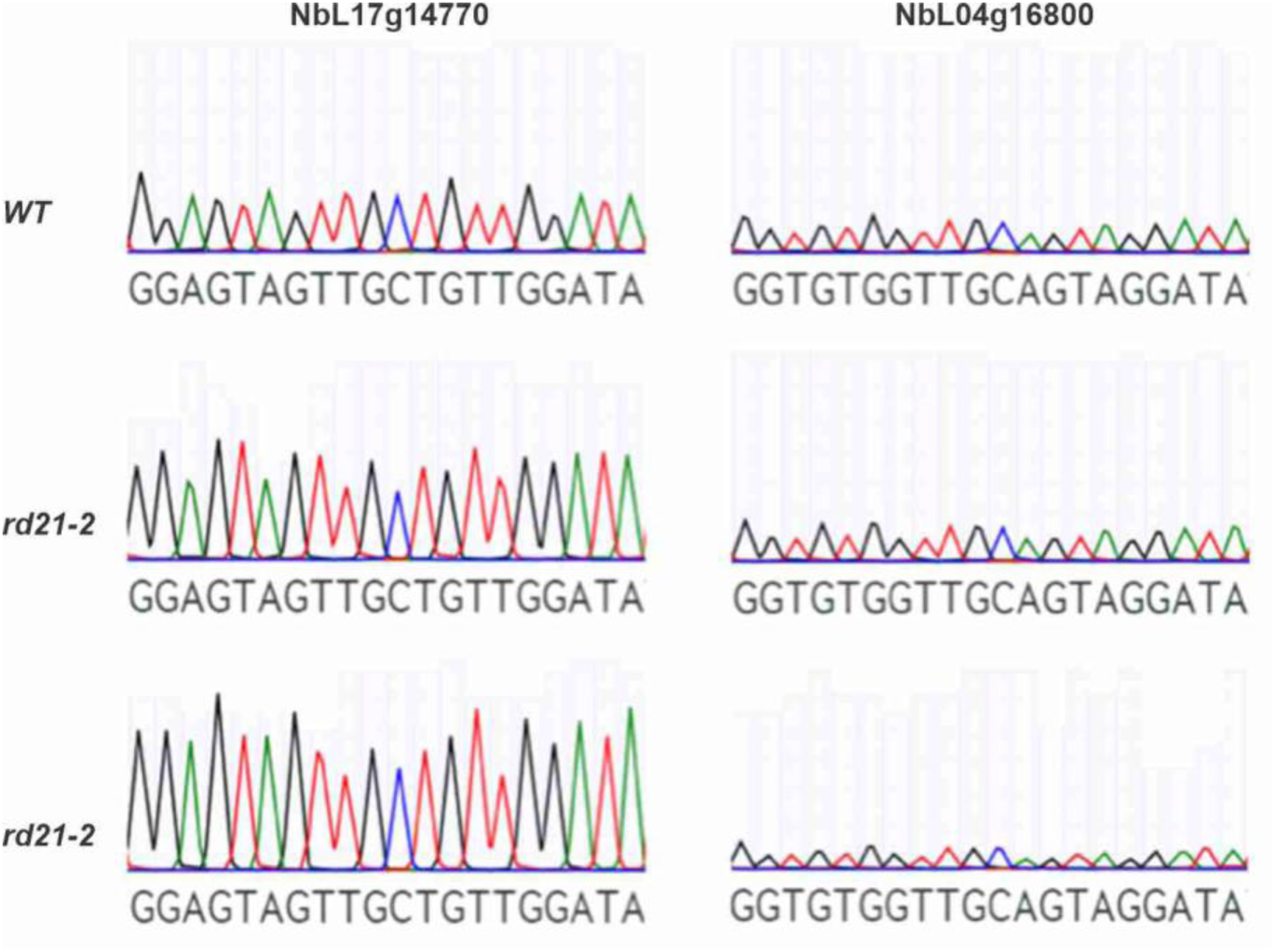
Unaltered off-targets of used sgRNAs. Cas-OFFinder identified sequences in the *N. benthamiana* genome with ≤3 mismatches with either sgRNA targeting NbL08g19980, returning RD21 homologs NbL17g14770 and NbL04g16800. Genes from wildtype (WT) and both *rd21* lines were sequenced and analysed using Benchling.com.

**Figure S3.**
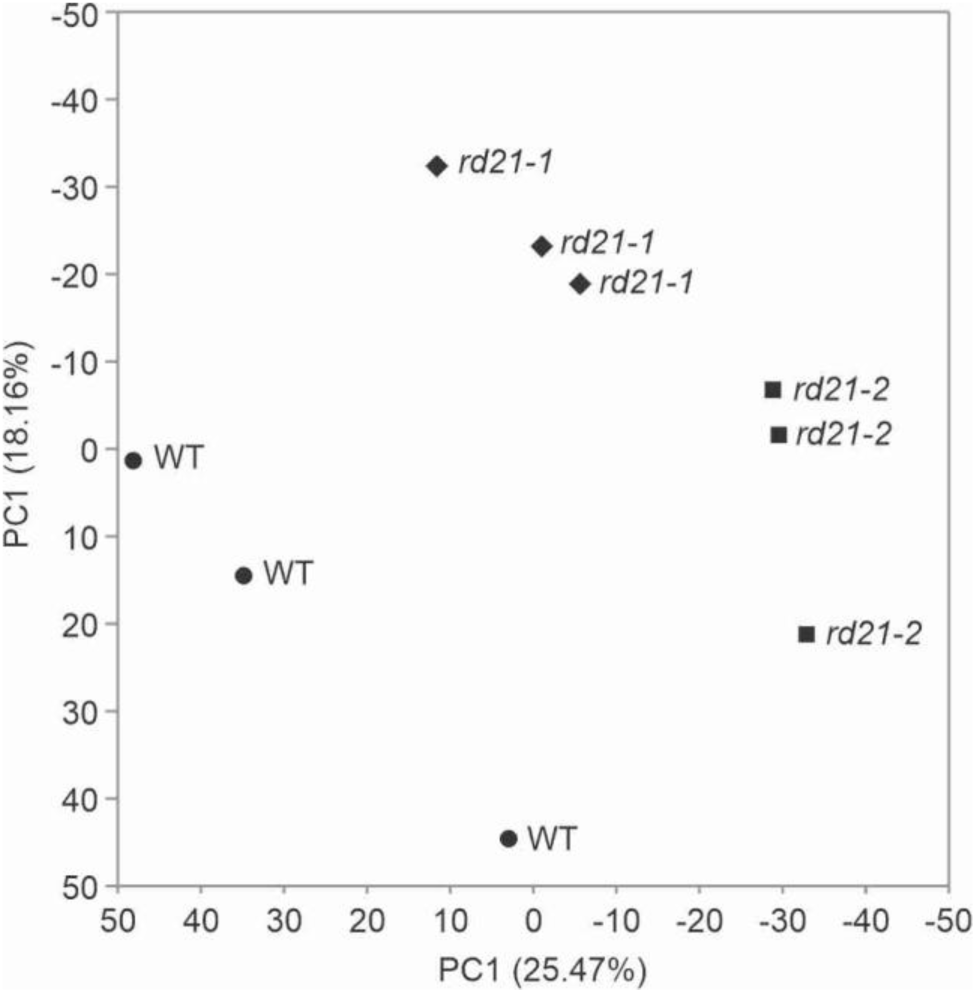
PCA analysis of the shot gun proteomics on the nine samples.

**Figure S4.**
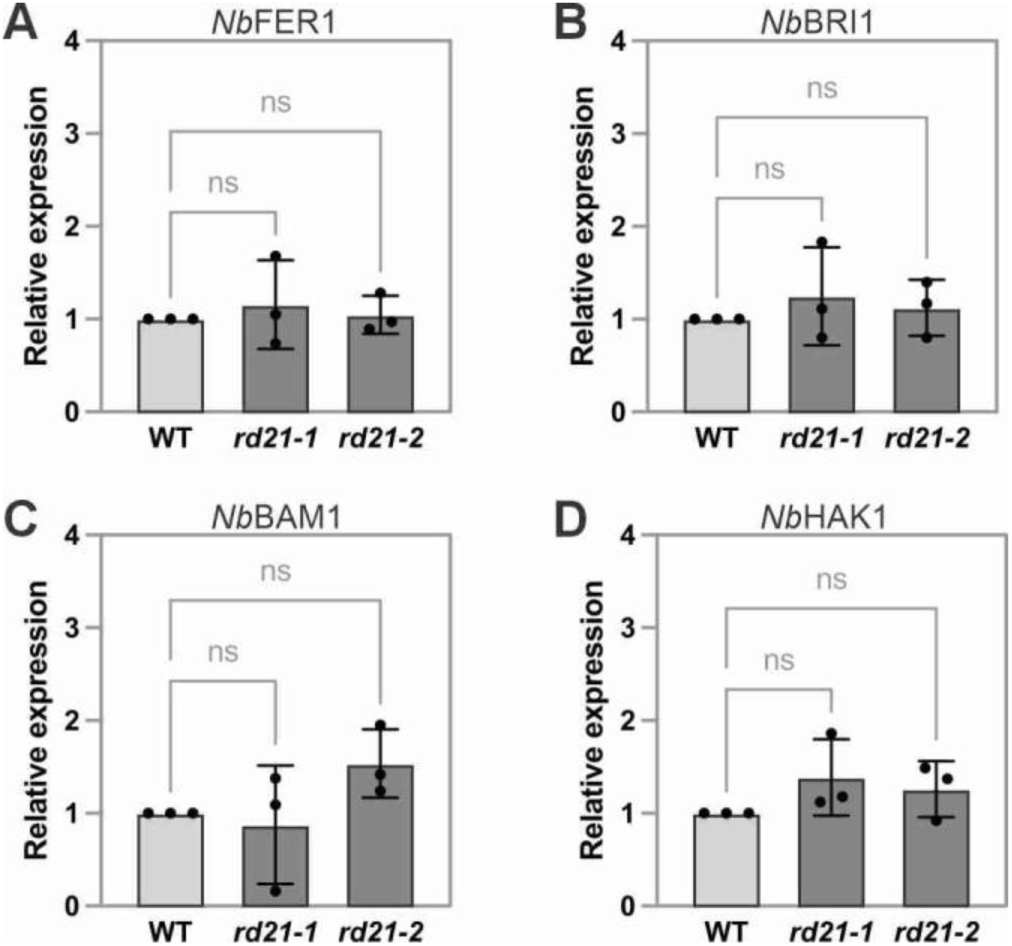
Transcript levels of RLK-encoding genes. Transcript levels were determined in n=3 plants of WT, *rd21-1* and *rd21-2* genotypes using gene specific primers, and *eIF4A* as a control, and normalized to WT plants. Error bars represent standard deviation of n=3 replicates. Statistical significance was evaluated using ANOVA plus Dunnett’s multiple comparison. ns, not significant.

**Figure S5.**
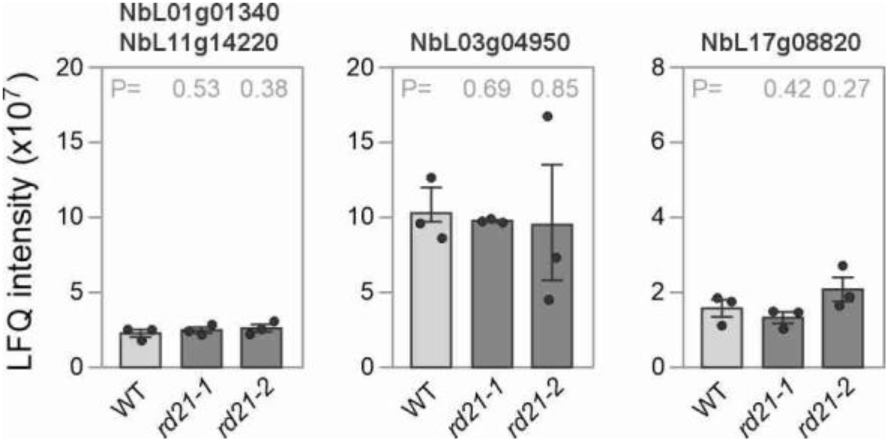
No increased accumulation of three RLKs in *rd21* plants Shown are raw LFQ intensities in wildtype (WT) and both knockout (*rd21-1* and *rd21-2*) plants. Error bars represent SE of n=3 samples. P-values calculated with Student’s *t*-test compared to WT were all insignificant.

**Figure S6.**
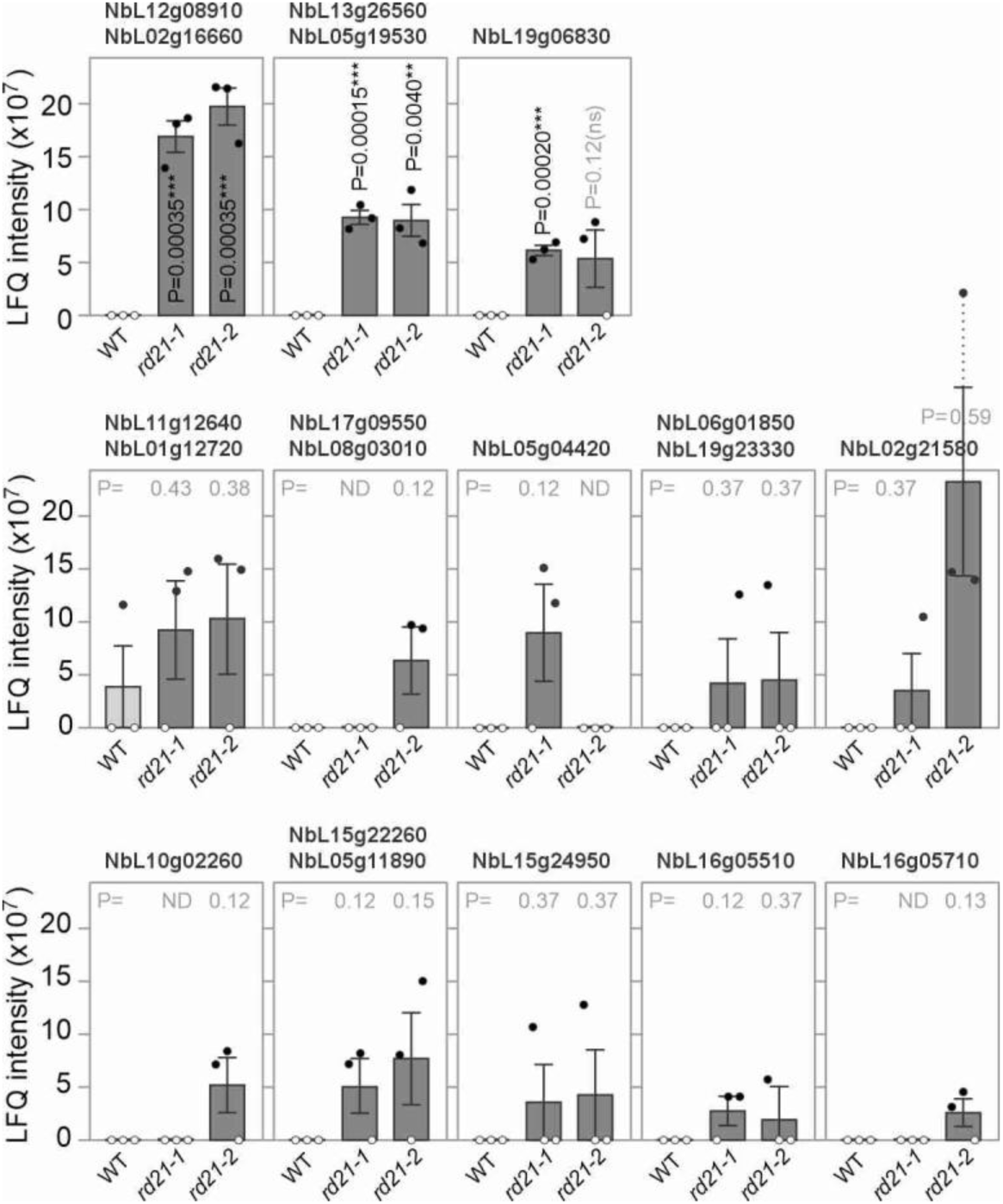
13 additional RLKs seem to accumulate more in *rd21* mutants. Bar charts showing raw LFQ intensities in wildtype (WT) and both knockout (*rd21-1* and *rd21-2*) plants. Open dots indicate the absence of a detectable MS signal. Error bars represent SE of n=3 samples. Student’s *t*-test compared to WT values: **, p<0.01; ***, p<0.001.

**Figure S7.**
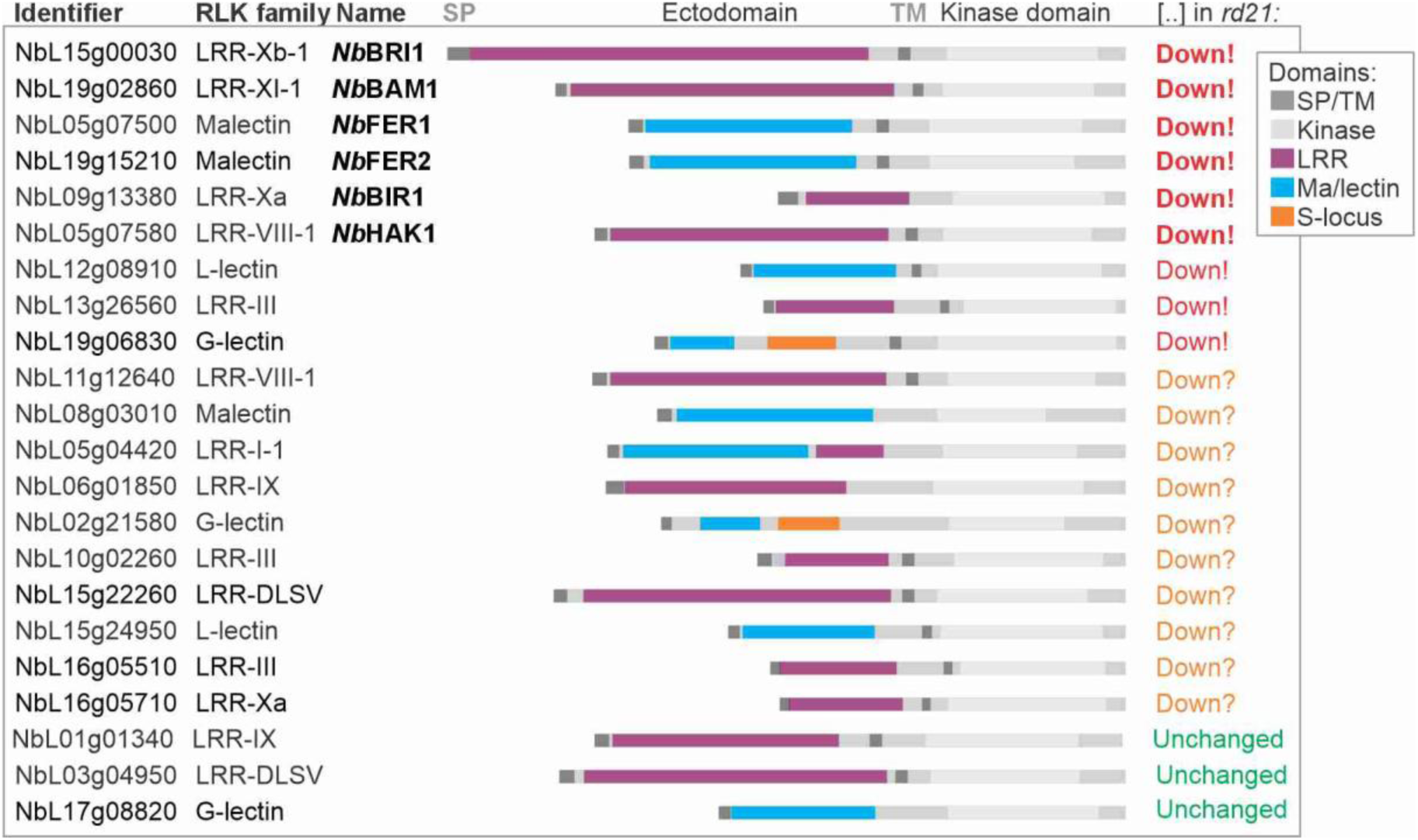
Domain structure of the detected RLKs Shown are the PFAM domains identified by InterPro and the signal peptide (SP) and transmembrane domains (TM), in addition to four types of ectodomains and the protein kinase domain. RLK families (second column) were assigned based on the metaRLK database (Zhang et al., 2026) and the homology to Arabidopsis LRR-RLKs. The RLKs are grouped based on their relative accumulation in *rd21* mutants (last column).

**Table S2.**
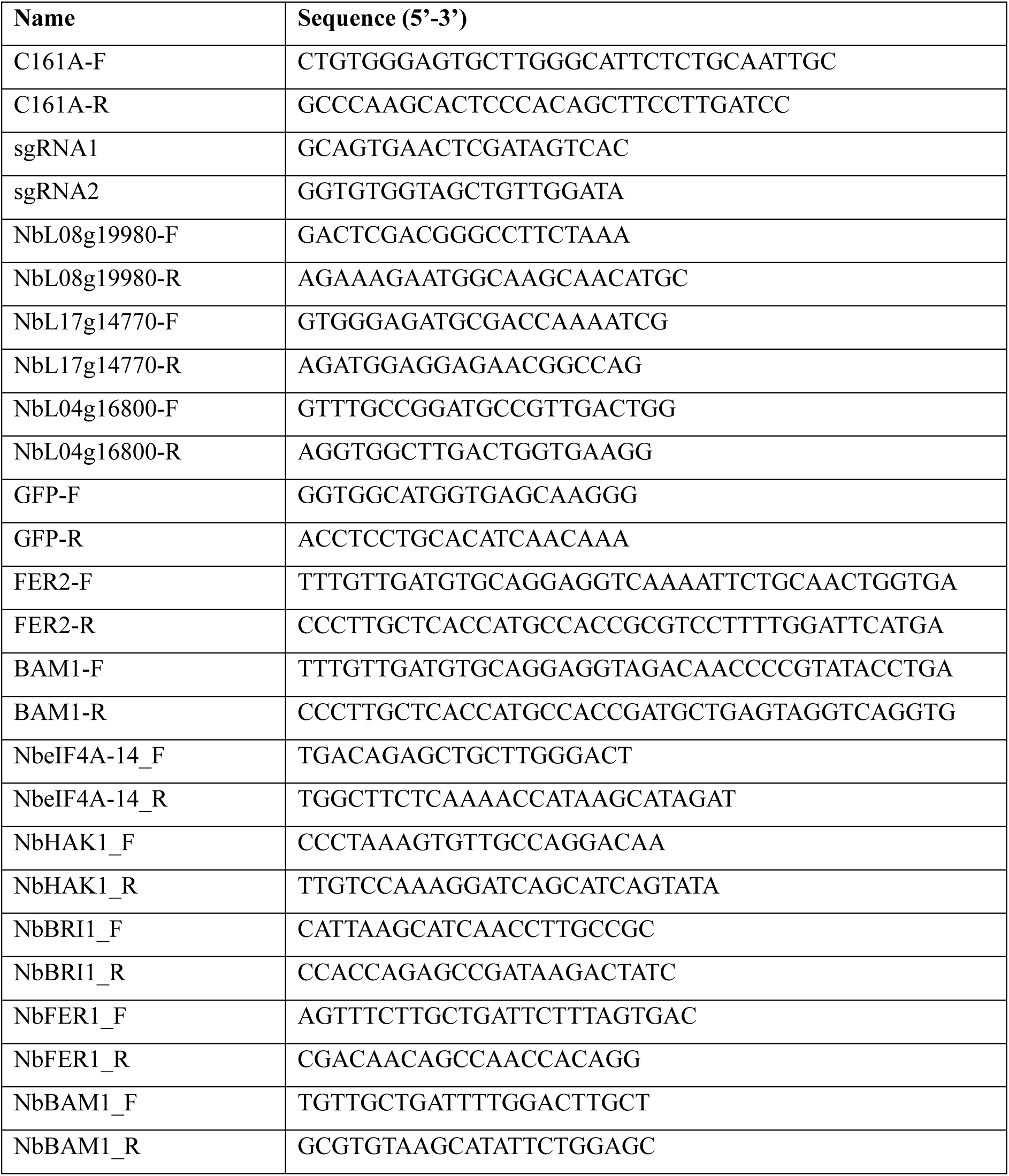
Used oligonucleotides.

**Table S3.** Used plasmids.

| Name | Description | Reference |
| --- | --- | --- |
| pJK001c | Empty vector | Paulus et al., 2020 |
| P19 | Binary 35S::P19 | Van der Hoorn et al., 2003 |
| pPB34 | Binary 35S::*Nb*RD21 | Sueldo et al., 2024 |
| pAG11 | Binary 35S::C161A | This work |
| pPJ064 | Binary RD21 genome editing plasmid | This work |
| pMS21 | Binary 35S::FER2-GFP | This work |
| pMS24 | Binary 35S::BAM1-GFP | This work |
| pJK227 | Binary 35S::Cf-4-GFP | Kourelis et al., 2020 |
| pJK309 | Binary Cas9 vector | This work |
| pID41 | Binary Cytoplasmic 35S::GFP (no p19) | Dodds et al., 2025 |
| pID42 | Binary Secreted 35S::RFP (no p19) | Dodds et al., 2025 |

